# Tuning of feedforward control enables stable muscle force-length dynamics after loss of autogenic proprioceptive feedback

**DOI:** 10.1101/851626

**Authors:** JC Gordon, NC Holt, AA Biewener, MA Daley

## Abstract

Animals must integrate feedforward, feedback and intrinsic mechanical control mechanisms to maintain stable locomotion. Recent studies of guinea fowl (*Numida meleagris*) revealed that the distal leg muscles rapidly modulate force and work output to minimize perturbations in uneven terrain. Here we probe the role of reflexes in the rapid perturbation response of muscle by studying the effects of proprioceptive loss. We induced bilateral loss of autogenic proprioception in the lateral gastrocnemius muscle (LG) using self-reinnervation. We compared ankle kinematics and *in vivo* muscle dynamics in birds with reinnervated LG and intact LG. Reinnervated and intact muscles exhibit similar force-length dynamics, with rapid changes in work to stabilize running obstacle terrain. Reinnervated LG exhibits 23ms earlier steady-state activation, consistent with feedforward tuning of activation phase to compensate for lost proprioception. Modulation of force duration is impaired in rLG, confirming the role of reflex feedback in regulating force duration in intact muscle.

## Introduction

Sensory feedback is widely accepted as an integral component of vertebrate locomotor control (Cohen, 1992; Donelan & Pearson, 2004; Grillner, 2011; Prochazka & Ellaway, 2012; Rossignol et al., 2006). The function of proprioceptive feedback can be conceptualized as two main components: 1) short-latency reflex feedback via mono- and polysynaptic pathways to regulate the ongoing activity and mechanical output of muscles (force, stiffness, impedance and work) and 2) longer-latency feedback pathways to spinal rhythm networks and higher CNS areas to estimate state and update internal models to coordinate balance and plan movement (Frigon & Rossignol, 2006; Proske & Gandevia, 2012; Rossignol et al., 2006; Wolpert, et al., 2011). Proprioception has been shown to regulate limb force during stance and limb cycling rhythm, including stance-swing and swing-stance transitions (Grillner, 2011; Lam & Pearson, 2002; Prochazka & Ellaway, 2012; Sherrington & Laslett, 1903; Sherrington 2010). These functions are fulfilled through autogenic (self-generated) monosynaptic reflexes to each muscle, and through poly-synaptic projections that modulate multiple muscles via spinal interneurons (Frigon & Rossignol, 2006; Lam & Pearson, 2002; Ross & Nichols 2009). Proprioception also contributes to inter-joint coordination through a heterogenic inter-muscular network of feedback (Abelew, Miller, Cope, & Nichols, 2000; Ross & Nichols 2009). Thus, the relationship between a specific sensory signal and its resulting effects is complex and dynamic.

Despite recognized functions of proprioception, the relative contribution of feedback control in high-speed locomotion remains unclear. Sensorimotor delay constrains how quickly an animal can sense and respond to a stimulus using feedback control (More & Donelan, 2018; More et al., 2010). The fastest possible feedback loop occurs through mono-synaptic reflexes, which involve a delay that increases in proportion to nerve transmission distance (More & Donelan, 2018; More et al., 2010). This reflex delay becomes a larger fraction of the stride cycle with increasing speed, limiting time available for reflex-mediated correction.

The challenges of long delays relative to stride cycle times likely necessitates greater reliance on feedforward control strategies at higher speeds. Here we use feedforward to refer to the contributions to motor output originating from higher brain centers and descending pathways that regulate the drive to rhythmic spinal networks (Frigon & Rossignol, 2006; Pearson, 2000; Yakovenko, et al. 2004). These pathways can generate the basic flexion and extension motor pattern when proprioceptive feedback is removed (e.g., Pearson et al., 2003; Sharp & Bekoff 2015). In reality, these descending networks act in concert with feedback, relying on multimodal and distributed sensory inputs to update state estimates and regulate rhythm generation (Cohen 1992; Pearson et al. 1999; Todorov 2004; Wagner & Smith 2008; Wolpert et al. 2011). Consequently, there is no true ‘pure’ feedfoward control within vertebrate systems. We use the term here as a pragmatic distinction between longer-latency ‘anticipatory’ control and shorter-latency ‘reactive’ control. *Feedforward* here refers to anticipatory ‘look-ahead’ control taking place over one or more complete stride cycles, and *feedback* refers to short latency reactive responses to perturbations.

Although feedforward networks normally act in concert with feedback, feedforward motor activation coupled to intrinsic muscle properties can be sufficient to produce stable gait, provided that central drive is well-matched to anticipated load (Yakovenko et al., 2004). Consistent with this, the lateral gastrocnemius (LG) of guinea fowl (*Numida meleagris*) rapidly absorbs energy in response to unexpected drop perturbations (Daley et al., 2009), which stabilizes high speed running without a reflex response. Such rapid perturbation responses arise from feedforward muscle activation coupled to intrinsic mechanical properties of the muscle-tendon tissues (Brown & Loeb 1999; Loeb et al. 1999). The intrinsic mechanical responses can be actively tuned by the specific feedforward timing and pattern of muscle activation. For example, humans hopping on surfaces with randomized sudden increases in ground stiffness show an anticipatory, feedforward increase in knee flexion and leg muscle co-activation, increasing mechanical stability (Moritz & Farley 2004). Many perturbations involve coordinated feedforward, feedback and mechanical contributions that overlap in time. Guinea fowl running over obstacles employ a combination of feedforward, intrinsic mechanical and reflex-mediated responses to obstacle encounters, with a sensorimotor delay of ~30-40 ms for reflex-mediated activity (Daley & Biewener, 2011; Gordon et al. 2015). Considering that the feedforward and intrinsic mechanical contributions alter the ongoing muscle dynamics *before* the reflex-mediated response, it is difficult to disentangle the specific contributions of proprioceptive reflexes to the rapidly stabilizing response observed in the muscle dynamics (Daley & Biewener, 2011; Gordon et al. 2015).

### Investigating the role of proprioception through self-reinnervation

Here we probe the integration of feedforward, feedback and intrinsic mechanical control by eliciting a proprioceptive deficit in the lateral gastrocnemius muscle (LG) of the guinea fowl (*Numida meleagris*) using bilateral self-reinnervation. Self-reinnervation involves peripheral nerve branch transection and immediate repair, resulting in recovery of motor output with long-term, local loss of autogenic muscle proprioception (Cope et al., 1994; Bullinger et al. 2011). Self-reinnervation occurs through axonal regrowth and reconnection with denervated tissues over a recovery period of 4-8 weeks (Carr, et al., 2010; Cope, et al., 1994; Gordon & Stein, 1982; Vannucci et al., 2019). Reinnervated muscles remain unresponsive to stretch due to synaptic retraction of muscle spindle afferents from spinal cord lamina IX, leading to disconnection from parent motoneuron populations (Alvarez et al., 2011). However, intermuscular force and length feedback networks may remain at least partially intact (Lyle et al. 2016). Cats with reinnervated muscles show normal muscle activity and inter-joint coordination during level, slow gait (Maas et al., 2007), with deficits in ankle-knee coordination on a decline (Abelew et al., 2000). They also exhibit high variance in inter-joint coordination, but preservation of whole-limb function (Chang et al. 2009). These findings highlight the ability of animals to flexibly exploit musculoskeletal plasticity to maintain function and suggest self-reinnervation as a promising tool to investigate sensorimotor control mechanisms.

Study of neuromuscular control in the guinea fowl, a bipedal animal model, provides insight into similarities and differences among vertebrates that may relate to locomotor modality, evolutionary history, or both. Birds share key features of sensorimotor structure and function with mammals, including muscle tissue properties (Nelson, et al., 2004; Poore et al., 1997) and muscle proprioception through muscle spindle and Golgi tendon organs (Dorward, 1970; Haiden & Awad 1981; Maier, 1992). Ground birds use bipedal walking and running gaits with mechanics, energetics and stability demands that are similar to human locomotion (Daley & Birn-Jeffery, 2018; Gatesy & Biewener, 1991; Heglund, et al., 1982; Taylor, et al., 1982). Bipedal gaits also limit the redundancy of control mechanisms and poses a challenge for dynamic stability. Whereas a quadrupedal cat or rat could compensate for deficits by shifting weight bearing among legs, a biped with a bilateral proprioceptive deficit cannot. Accordingly, we expect the response to proprioceptive deficit to be more constrained in a biped compared to a quadruped. Additionally, bipedal gait involves substantial periods of single-limb contact, which simplifies biomechanical analysis and facilitates an integrative understanding of muscle, limb and whole-body dynamics (Roberts, et al., 1997; Nishikawa et al., 2007; Daley, et al., 2009; Clark & Higham, 2011; Daley & Biewener, 2011).

We hypothesize that autogenic proprioceptive deficit will lead to increased reliance on feedforward control mechanisms and intrinsic muscle mechanics to maintain stable locomotion. To test for shifts in stability and control mechanisms, we measured ankle kinematics and *in vivo* LG muscle dynamics (length, force and activation) during treadmill running on level and obstacle terrain. There are several possible mechanisms that could potentially compensate for autogenic proprioceptive deficit: 1) If feedback regulation of LG activity is essential for stability during fast locomotion, loss of LG autogenic proprioception may necessitate increased heterogenic reflex gains from synergists, with a slight delay compared to intact animals, as observed in cats (Pearson et al. 1999). 2) Alternatively, birds with proprioceptive deficit might compensate through increased feedforward muscle activation before obstacle contact, as observed in birds negotiating high-contrast, visible obstacles (Gordon et al. 2015). 3) Finally, if intrinsic mechanics are mainly responsible for the stabilizing modulation of muscle force and work, we might expect minimal change in muscle activity patterns (EMG). Considering the stability demands and constrained musculoskeletal dynamics of fast bipedal locomotion, we expect birds with reinnervated LG to compensate for proprioceptive deficit by tuning the feedforward muscle activity to maintain a stable response to obstacle perturbations. If stability is substantially impaired following reinnervation, this should be evident from altered kinematics, increased variance and longer time to recover from obstacle perturbations. By investigating the specific shifts in guinea fowl LG muscle force, length and activation dynamics following reinnervation, we hope to gain insight into the mechanisms of sensorimotor integration and plasticity that enable robustly stable and agile bipedal locomotion

## Methods

### Animals and training

We obtained and reared six hatchling guinea fowl keets (*Numida meleagris*) from a breeder (Hidden Hollow Acres, Whitehouse Station, NJ), to allow re-innervation surgeries in juveniles with at least 12 weeks for recovery before *in vivo* muscle procedures (see below). At the time of the *in vivo* muscle measurements, the guinea fowl had reached adult size, averaging 1.84±0.26 kg body mass. Birds had primary feathers clipped and were trained to run on a level motorized treadmill (Woodway, Waukesha, WI, USA) at speeds ranging from 0.5-5.0 ms^−1^. Training sessions were 15-20 minutes in duration, with breaks for 2 minutes as needed. We trained all subjects for 3 days per week for 1-2 weeks before data collection. All experiments were undertaken at the Concord Field Station of Harvard University, in Boston (MA, USA), and all procedures were licensed and approved by the Harvard Institutional Animal Care and Use Committee (AEP #20-09) in accordance with the guidelines of the National Institutes of Health and the regulations of the United States Department of Agriculture.

### Anesthesia and post-operative care

Birds were induced and maintained on a mid-plane of anesthesia using isoflurane (2 - 3%, mask delivery). We administered perioperative enrofloxacin and flunixin intramuscularly for analgesia after induction and continued for three days after each surgery. Birds recovered to bilateral weight bearing within 20 minutes following completion of surgical procedures.

### Reinnervation surgery

The timing of surgeries was planned based on a pilot study, which found full recovery of LG motor activity by 6 weeks following reinnervation surgeries, and continued absence of calcaneal tendon reflex one year later, indicating continued absence of autogenic stretch reflexes (Carr et al., 2010). We bilaterally transected and immediately repaired the peripheral nerve branch supplying the LG muscle in maturing guinea fowl between 7- 12 weeks of age. We then allowed time for full reinnervation recovery of motor output and growth to adult size before a subsequent surgery to implant muscle transducers (Fig. 1). In the reinnervation surgery, a lateral incision was made posterior-distal to the knee to expose the underlying muscle. Blunt dissection enabled exposure and identification of relevant nerve branches, and the identity of the correct nerve branch was confirmed using an isolated nerve stimulator (SD48, Grass Instruments, Warwick, RI) to visualize contraction in the LG. After pre-placement of single longitudinal throw of 6-0 braided non-absorbable silk (Silk, Ethicon, Somerville, NJ, USA) through a 3 mm nerve section, we transected the nerve branch and sutured to appose the cut nerve endings. Fibrin glue (bovine thrombin in CaCl_2_, fibrinogen, fibronectin from bovine plasma) was applied over the apposed nerve endings as an additional repair scaffold (Carr et al., 2010; Spotnitz, 2010). We closed the fascia and skin with 3-0 braided absorbable polyglactin (Vicryl, Ethicon, Somerville, NJ, USA).

**Figure 1.**
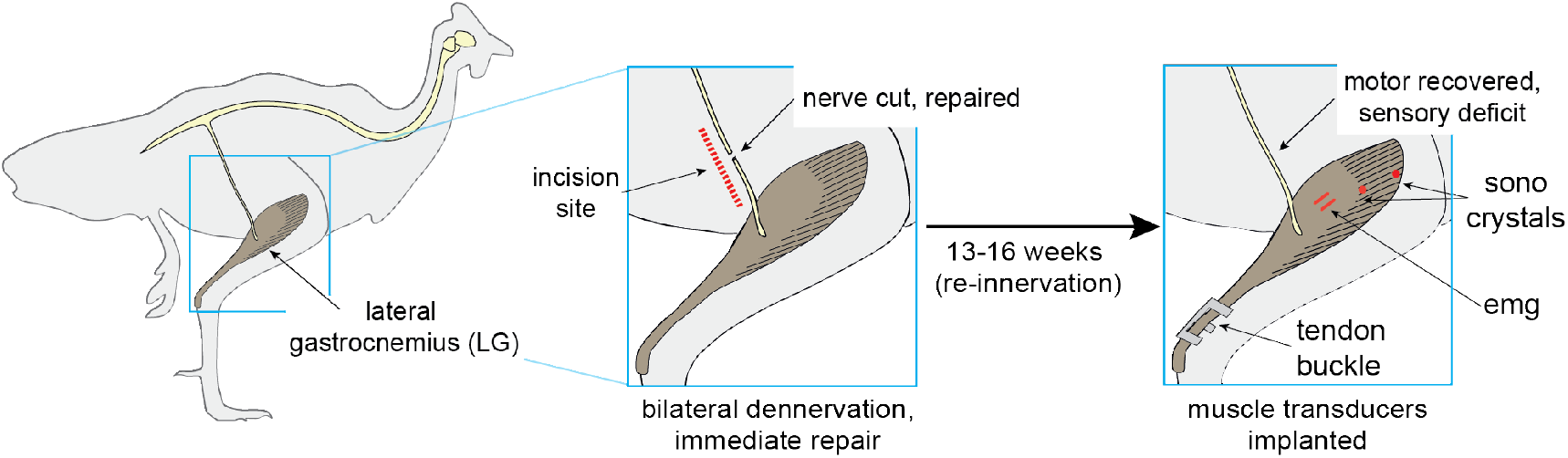
Protocol for bilateral self-reinnervation of the lateral gastrocnemius, followed by transducer implantation for *in vivo* muscle recordings.

In the immediate post-operative period, bilateral limb posture was visibly more crouched compared to ‘intact’ birds and Achilles tendon tap revealed no functional stretch reflex. LG atrophy was qualitatively observed during the first 2 weeks of the recovery period. Within 1 week of surgery, bird activity levels appeared comparable to intact conspecifics, with limb posture partially recovered. From 2-3 weeks onwards, differentiating reinnervated from ‘intact’ birds was not possible from grossly observable limb morphology, posture and gait. We conducted regular treadmill training from 7 weeks after reinnervation surgery. During training and experimental recordings, birds did not stumble or fall with noticeably greater frequency than observed in intact birds (Daley & Biewener 2011) and were able to maintain treadmill position over a similar speed range.

### Transducer implantation surgery

When the birds were 23-28 weeks old (13-16 weeks following bilateral reinnervation surgeries), we performed a second surgery for transducers placement, following similar procedures as Daley and Biewener (2003). The surgical field was plucked of feathers and gently cleaned with antiseptic solution (Prepodyne, West Argo, Kansas City, MO, USA). We tunneled transducer leads subcutaneously from a 1–2 cm incision over the synsacrum to a second 4–5 cm incision over the lateral left shank. Sonomicrometry crystals (2.0 mm; Sonometrics Inc., London, Canada) were implanted into the lateral head of the gastrocnemius (LG) along the fascicle axis in the middle 1/3rd of the muscle belly. Crystals were placed in small openings using fine forceps, approximately 3–4 mm deep and 15 mm apart. We verified signal quality using an oscilloscope and secured the crystals by closing the overlying muscle fascia and lead wires with separate 4-0 silk sutures (Silk, Ethicon, Somerville, NJ, USA). Next to the crystal pair, we implanted bipolar EMG electrodes constructed from two strands of 38-gauge Teflon-coated stainless steel (AS 632, Cooner Wire Co., California, USA) with staggered 1 mm exposed regions spaced 1.5 mm apart. Electrodes were placed using sew-through methods and surface silicon anchors (3 × 3 × 2 mm) positioned with a single square knot at the muscle surface-electrode interface (Deban & Carrier, 2002). An “E”-type stainless-steel tendon buckle force transducer insulated with a polyurethane coating (Micro-Measurements, Raleigh NC) was implanted on the common gastrocnemius tendon, equipped with a metal foil strain gauge (type FLA-1, Tokyo Sokki Kenkyujo). We connected transducers to a micro-connector plug (15-way Micro-D, Farnell Ltd, Leeds, UK) sutured to the bird’s dorsal synsacrum.

### Transducer recordings

A lightweight shielded cable was used to connect the microconnector to data acquisition systems. Sonomicrometry data were collected via a Sonometrics TRX analog data-acquisition device and PC interface (TRX Series 8, Sonometrics, Ontario, Canada). Crystals were tested before surgery in a saline bath to confirm distances measured by digital caliper matched those measured by the software. Occasional level-shift artifacts (instantaneous length change errors due to variation in signal characteristics) were removed using a custom script in MATLAB (Mathworks, Inc.; Natick, MA, USA). Tendon buckle signals were fed through a bridge amplifier (Vishay 2120, Micro-Measurements, Raleigh, NC), and EMG signals were amplified and bandpass filtered (10Hz and 3kHz) using GRASS pre-amplifiers (P511, Grass Instruments, Warwick, RI). Signals were recorded at 5000 Hz using a 16-channel, 16-bit Biopac A/D acquisition device (MP150, BIOPAC systems, Gotleta, CA, USA).

Following experiments, birds were euthanized using an intravenous injection of sodium pentobarbital (100 mg kg^−1^) while under deep isoflurane anesthesia (4%, mask delivery). We recorded the morphology of the muscle and the location of transducers to confirm muscle fascicle and tendon alignment. LG mass was 10.0±2.0 g, and the medial gastrocnemius (MG) was 15.4±3.7 g. These muscle masses are a comparable to those measured from intact LG and MG in previous studies, representing approximately 0.5% and 0.8% body mass for LG and MG, respectively (Daley & Biewener 2003; Higham & Biewener 2008). This suggests recovery from prior denervation-induced muscle atrophy. Fascicle lengths were 17.6±3.1 mm and 21.1±4.0 mm, tendon lengths 45.2±16.0 mm and 53.4 ± 16.6 mm and pennation angles 23.8 ± 5.2º and 16.2 ± 4.2º, for LG and MG, respectively. Crystal alignment relative to the fascicle axis (*α*) was within 2°, indicating that errors due to misalignment were <0.1%. We calibrated the tendon force buckle *in situ post mortem* by applying a series of known cyclical loads using a force transducer (model 9203, Kistler, Amherst, MA), which yielded linear least-squares calibration slopes with R^2^ > 0.97.

### Experimental protocol

The guinea fowl were run at 1.7-2.0 ms^−1^ for 30-second trials spaced with 10-minute rest periods with access to food and water. We recorded trials at the same speed for uniform level terrain and terrain with 5 cm obstacles. The speeds and terrain conditions were consistent with those recorded for intact guinea fowl (Daley & Biewener 2011), to allow statistical comparison of the two groups (n=6 each for reinnervated and intact birds).

The treadmill belt (Woodway, Waukesha WI) was slatted black rubber-coated steel with running surface 55.8 cm × 172.7 cm with clearance for obstacles beneath. Obstacles were constructed from styrofoam reinforced with cardboard covered with black neoprene to form a light, stiff surface. Waterproof glue (Shoe Goo, Eugene, OR, USA) secured heavy-duty fabric hook and loop fastener (Velcro, Cheshire, UK) to the obstacle and treadmill surface. Four sequential slats of obstacles produced a 20 cm^2^ continuous obstacle surface. Obstacles were encountered approximately every 4-5 strides, with some variation due to varied stride length and treadmill station keeping. We recorded high-speed video at 250 Hz (Photron, San Diego, CA, USA) for analysis of ankle kinematics, detection of stride timing and statistical coding of strides in relation to obstacle contact.

### Data Processing

The analyzed data includes all strides in the level and obstacle terrain. In total, we analyzed 1024 strides for reinnervated individuals (n=6) and 1511 strides for intact individuals (n=6) across level and obstacle terrain conditions. Data from the intact individuals was published previously in Daley and Biewener (2011) and is included here to allow for direct statistical comparison. The dataset analyzed here includes a larger sample of strides and more variance in obstacle contact conditions than reported in Daley and Biewener (2011), because the previous analysis included only the obstacle strides in which the foot stepped cleanly and directly onto the obstacle with no previous brushing of the foot or toes.

We assigned strides categories in relation to the obstacle encounters, using the same approach as in Daley and Biewener (2011), which considers the stride sequence of the instrumented leg: the stride prior to an obstacle contact (S −1), obstacle contact strides (S 0), the stride following obstacle contact (S +1), and strides within the level region between obstacles (S +2). All level terrain strides were assigned the same stride category (L). Note, this is simpler stride coding than presented in Gordon and colleagues (2015), which also considered the timing of obstacle encounters by the contralateral leg. Strides in which the contralateral limb had made contact with the obstacle in the previous step are grouped here into the (S +2) category, for simplicity. This does not substantially alter the findings, because the coding was similar between intact and reinnervated birds, and the analysis here is focused on understanding the shifts in LG force-length dynamics in relation to a direct obstacle encounter by the instrumented leg.

Features of LG activation, force-length dynamics and work output were measured, similar to Daley and Biewener (2011). Raw EMG signals were used to calculate myoelectric intensity in time and frequency domains using wavelet decomposition (Daley et al. 2009; Gordon et al. 2015). This was used to calculate total myoelectric intensity per stride (E_tot_) and mean frequency of muscle activation (E_freq_). We calculated fractional fascicle length (L) from sonomicrometry data using mean length in level terrain as a reference length (L_o_). Note, however, that L_o_ is not directly related to sarcomere length or optimal length for the isometric force-length curve, which were not measured. Fractional fascicle length was differentiated to obtain fascicle velocity (V, in lengths per second, Ls^−1^). Shortening strains are negative. We multiplied fascicle velocity (in ms^−1^) by tendon force (in Newtons, N) to calculate muscle power (Watts), which was integrated through time to calculate total work per stride (Joules, J; with shortening work being positive), and then normalized by muscle mass to obtain mass-specific muscle work (Jkg^−1^). We also recorded muscle (fascicle) length, velocity and force at specified times to evaluate how strain and activation factors influence muscle force and work output. All data processing was completed using MATLAB (Mathworks, Inc.; Natick, MA, USA).

### Statistics

We used a linear mixed-effects model ANOVA with treatment (intact/reinnervated) and stride category (stride ID: S −1, S 0, S +1, S +2, L) as fixed categorical factors, including the fixed-effects interaction term (treatment x stride ID), and individual as a random effect. This model was found to have a lower Akaike Information Criterion (AIC, Akaike, 1976) than comparison models with subsets of these factors. Statistical analysis was completed in Matlab using ‘fitlme’ and associated functions in the Statistics and Machine Learning Toolbox (Mathworks, Inc.; Natick, MA, USA). Post-hoc pairwise comparisons are based on the 95% confidence intervals for the coefficients from the linear mixed effects model.

## Results

We find that the general pattern of *in vivo* force-length dynamics of the guinea fowl reinnervated lateral gastrocnemius (rLG) is qualitatively similar to that previously measured in intact birds. An illustrative ten-stride sequence of length, force and EMG recordings is shown for rLG in Figure 2. During the swing phase of the stride cycle, rLG exhibits period of passive stretch, followed by shortening that begins shortly after mid-swing. Activation and force development begin in late swing around the time of the transition from stretch to shortening, with activation initiating rapid active shortening until foot contact (Fig. 2, toe down at dashed vertical lines, with light grey bird silhouette). At the time of foot contact, force increases rapidly to peak before midstance and then declines more slowly. Typically, rLG shortens slowly throughout force development, producing net positive work. However, muscle fascicle length change during contraction can vary between a stretch-shorten trajectory in some strides (e.g., Fig. 2, stride 3) and continuous shortening in others (e.g., Fig. 2, stride 5).

**Figure 2.**
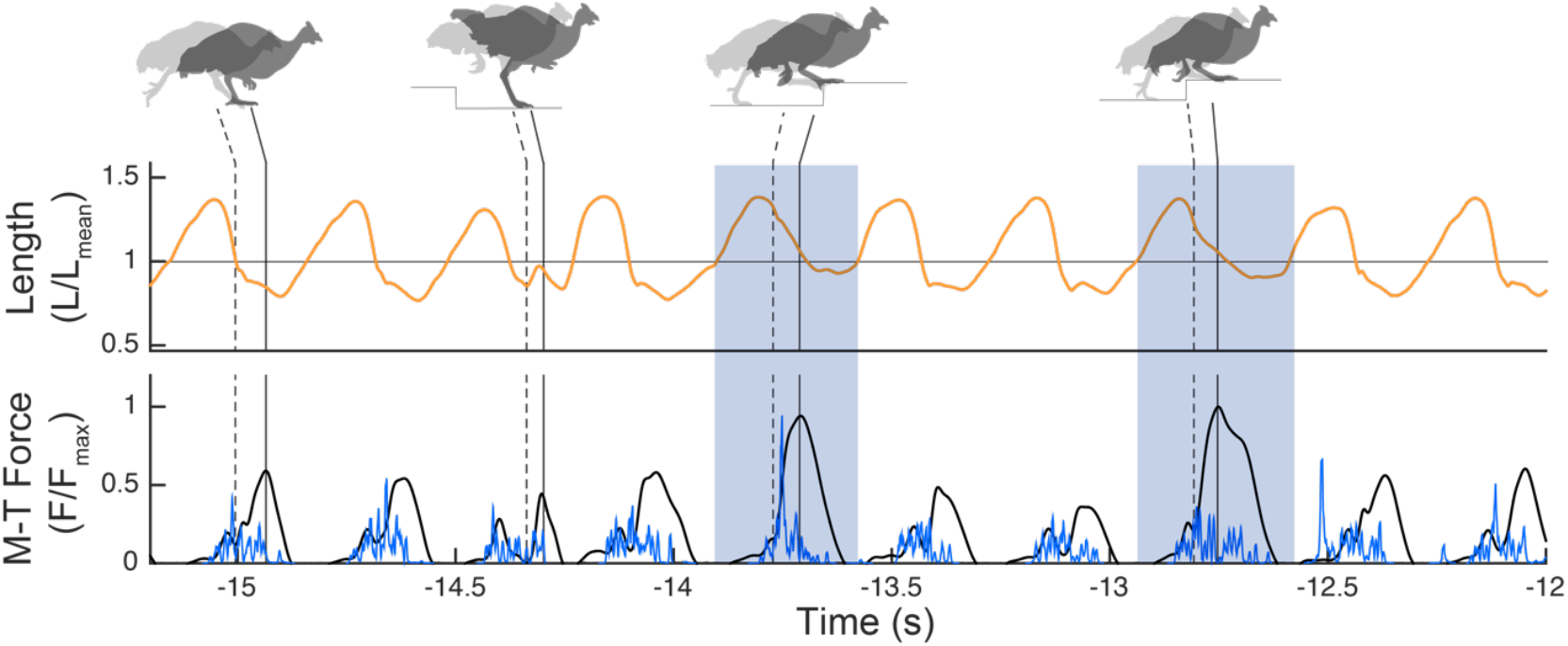
Example 10-stride sequence of *in vivo* muscle recordings of the reinnervated lateral gastrocnemius (rLG) in the right leg, running at 1.7ms^−1^ on the obstacle treadmill. Muscle length (top, orange), force (bottom, black) and activation (bottom, blue) are shown, with a shaded box indicating obstacle encounter strides (S 0). Silhouettes at the top illustrate the leg posture at the time of foot contact light grey silhouette, dashed vertical line indicates timing) and at the time of peak muscle-tendon force (dark grey silhouette, solid vertical line indicates timing), taken from video. Note that between stride 2 and 3 the contralateral leg stepped on the obstacle, leading to a downward step of the instrumented leg in stride 3.

### Force-length dynamics during obstacle negotiation

The guinea fowl LG exhibits rapid increases in force and work output upon foot contact with an obstacle, and this stabilizing response remains qualitatively similar between intact and reinnervated conditions (Fig. 3). Upon obstacle contact, LG remains at longer lengths, reaches higher peak force and shortens steadily throughout force development (Fig. 3, S 0). This corresponds to an increased total impulse (integral of force) and net positive work produced during the stride cycle (the area enclosed within the work loop). In S 0, the net total work produced (W_net_) increases by 4.23 ± 2.03Jkg^−1^ (mean±S.D) in intact LG (iLG) and 4.44±2.44Jkg^−1^ in rLG compared to steady-state level strides. In Figure 4, average length, force and EMG trajectories are shown over time for example individuals with intact and reinnervated LG, for both level and obstacle terrain conditions. Notably, despite deficits in LG monosynaptic reflex following reinnervation, rLG and iLG show similar increases in total muscle activation intensity (E_tot_, integral of EMG) in obstacle strides (S 0, Fig. 4, Table 1 & 2). The timing and mechanisms of the increased activation will be explored further below.

**Figure 3.**
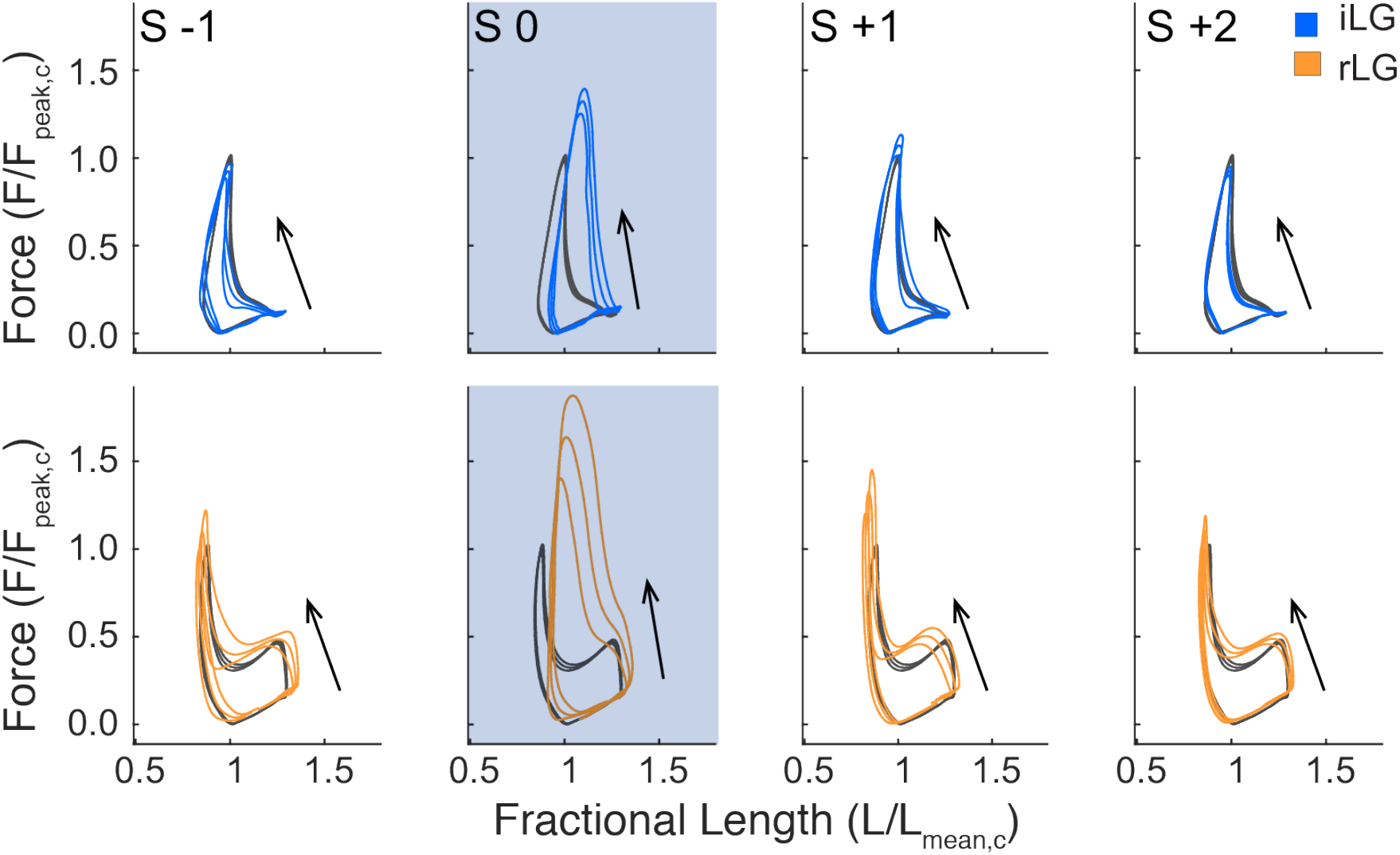
Force-length work loops for an intact (iLG) and reinnervated LG (rLG), for a single individual from each condition (mean ± 95% confidence intervals). Level terrain strides in grey and obstacle terrain strides in colored lines (top: blue, intact LG, bottom: orange, reinnervated LG). Stride categories are coded as in Daley and Biewener 2011 according the obstacle negotiation sequence, preceding (S −1), on (S 0) and following obstacle contact (S +1, S +2), with the shaded box corresponding to an obstacle encounter with the instrumented leg.

**Figure 4.**
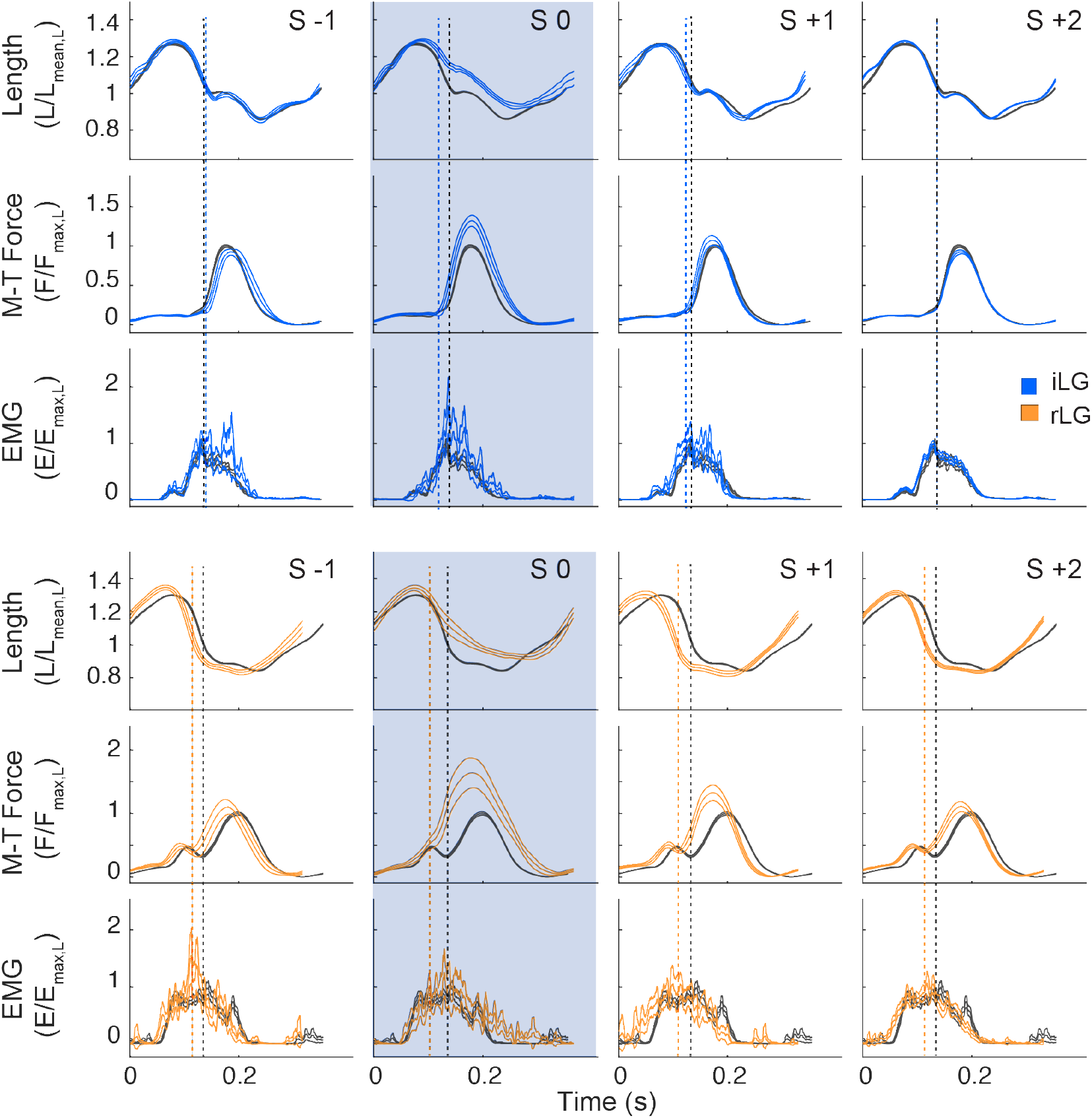
Comparison of intact (iLG, blue, top) and reinnervated (rLG, orange, bottom) average stride cycle trajectories (mean± 95% confidence intervals) for fascicle length, muscle-tendon force and myoelectric activity (EMG) during obstacle negotiation. Level terrain averages shown in grey. Example data are from one individual in each treatment category (blue: iLG, orange: rLG). Vertical dashed lines indicate the timing of foot contact with the substrate (black: level terrain, color: obstacle terrain). The shaded box highlights the obst
acle contact stride (S 0).

**Table 1.** F-statistics for linear mixed effect model ANOVA on measures of muscle contraction dynamics. Bolding indicates statistical significance (P <= 0.05). Individual was included as a random factor. Degrees of freedom for fixed effects were Treatment = 1, StrideID = 4, Interaction (Treatment × StrideID) = 4, and Error = 2525.

| Variable | F-statistic |  |  |
| --- | --- | --- | --- |
|  | Treatment | StrideID | interaction |
| $W_{net}$ | 3.77 | **179.19** | **7.52** |
| $W_{load}$ | 0.04 | **89.34** | **24.65** |
| $F_{pk}$ | 0.50 | **29.93** | **60.00** |
| $J_{tot}$ | 0.11 | **108.24** | **46.84** |
| $L_{T50}$ | **6.01** | **255.97** | **31.42** |
| $L_{pkF}$ | 1.50 | **234.48** | **41.55** |
| $V_{mean}$ | **11.32** | **88.23** | **23.79** |
| $V_{pkF}$ | **11.48** | **28.03** | **8.81** |
| Force duration | 2.46 | **160.86** | **28.36** |
| Force rate | 0.45 | **14.57** | **21.86** |
| Stride duration | 0.10 | **12.40** | **18.02** |
| $E_{tot}$ | 0.00 | **50.70** | **3.74** |
| $E_{freq}$ | **3.94** | **8.31** | 1.95 |
| $E_{phase}$ | **5.83** | 2.15 | **8.60** |

**Table 2.**
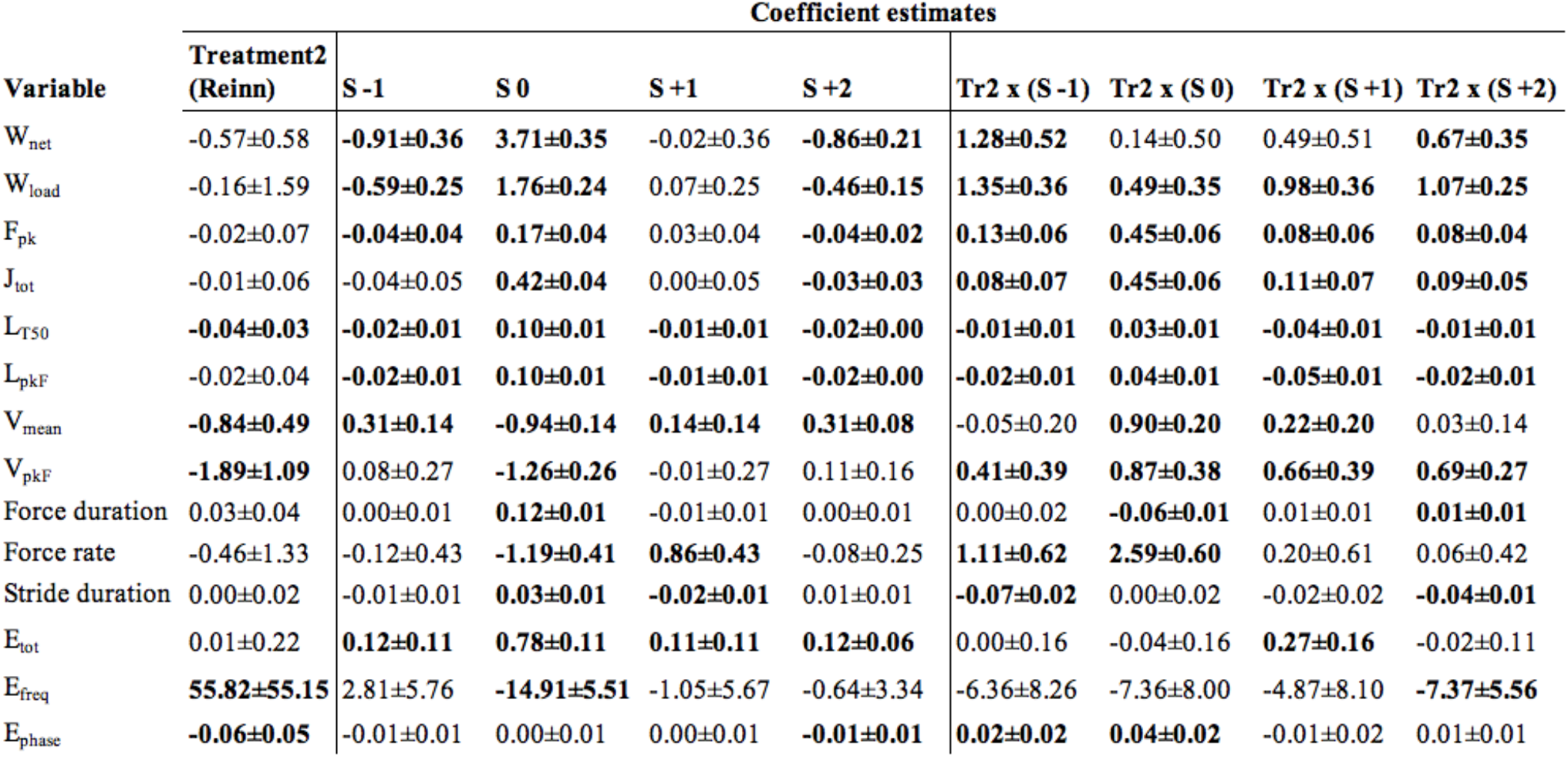
Fixed effect coefficients for the linear mixed effects model (estimate + 95% confidence interval).

| Variable | Coefficient estimates |  |  |  |  |  |  |  |  |
| --- | --- | --- | --- | --- | --- | --- | --- | --- | --- |
|  | Treatment2<br>(Reinn) | S -1 | S 0 | S +1 | S +2 | Tr2 x (S -1) | Tr2 x (S 0) | Tr2 x (S +1) | Tr2 x (S +2) |
| $W_{net}$ | -0.57±0.58 | **-0.91±0.36** | **3.71±0.35** | -0.02±0.36 | **-0.86±0.21** | **1.28±0.52** | 0.14±0.50 | 0.49±0.51 | **0.67±0.35** |
| $W_{load}$ | -0.16±1.59 | **-0.59±0.25** | **1.76±0.24** | 0.07±0.25 | **-0.46±0.15** | **1.35±0.36** | **0.49±0.35** | **0.98±0.36** | **1.07±0.25** |
| $F_{pk}$ | -0.02±0.07 | **-0.04±0.04** | **0.17±0.04** | 0.03±0.04 | **-0.04±0.02** | **0.13±0.06** | **0.45±0.06** | **0.08±0.06** | **0.08±0.04** |
| $J_{tot}$ | -0.01±0.06 | -0.04±0.05 | **0.42±0.04** | 0.00±0.05 | **-0.03±0.03** | **0.08±0.07** | **0.45±0.06** | **0.11±0.07** | **0.09±0.05** |
| $L_{T50}$ | **-0.04±0.03** | **-0.02±0.01** | **0.10±0.01** | **-0.01±0.01** | **-0.02±0.00** | **-0.01±0.01** | **0.03±0.01** | **-0.04±0.01** | **-0.01±0.01** |
| $L_{pkF}$ | -0.02±0.04 | **-0.02±0.01** | **0.10±0.01** | **-0.01±0.01** | **-0.02±0.00** | **-0.02±0.01** | **0.04±0.01** | **-0.05±0.01** | **-0.02±0.01** |
| $V_{mean}$ | **-0.84±0.49** | **0.31±0.14** | **-0.94±0.14** | **0.14±0.14** | **0.31±0.08** | -0.05±0.20 | **0.90±0.20** | **0.22±0.20** | 0.03±0.14 |
| $V_{pkF}$ | **-1.89±1.09** | 0.08±0.27 | **-1.26±0.26** | -0.01±0.27 | 0.11±0.16 | **0.41±0.39** | **0.87±0.38** | **0.66±0.39** | **0.69±0.27** |
| Force duration | 0.03±0.04 | 0.00±0.01 | **0.12±0.01** | -0.01±0.01 | 0.00±0.01 | 0.00±0.02 | **-0.06±0.01** | 0.01±0.01 | **0.01±0.01** |
| Force rate | -0.46±1.33 | -0.12±0.43 | **-1.19±0.41** | **0.86±0.43** | -0.08±0.25 | **1.11±0.62** | **2.59±0.60** | 0.20±0.61 | 0.06±0.42 |
| Stride duration | 0.00±0.02 | -0.01±0.01 | **0.03±0.01** | **-0.02±0.01** | 0.01±0.01 | **-0.07±0.02** | 0.00±0.02 | -0.02±0.02 | **-0.04±0.01** |
| $E_{tot}$ | 0.01±0.22 | **0.12±0.11** | **0.78±0.11** | **0.11±0.11** | **0.12±0.06** | 0.00±0.16 | -0.04±0.16 | **0.27±0.16** | -0.02±0.11 |
| $E_{freq}$ | **55.82±55.15** | 2.81±5.76 | **-14.91±5.51** | -1.05±5.67 | -0.64±3.34 | -6.36±8.26 | -7.36±8.00 | -4.87±8.10 | **-7.37±5.56** |
| $E_{phase}$ | **-0.06±0.05** | -0.01±0.01 | 0.00±0.01 | 0.00±0.01 | **-0.01±0.01** | **0.02±0.02** | **0.04±0.02** | -0.01±0.02 | 0.01±0.01 |

### Shifts in steady state force-length and activation dynamics between intact and reinnervated LG

Although the force-length dynamics of iLG and rLG are qualitatively similar, they exhibit several differences in specific features of muscle dynamics (Fig. 5). The rLG shows higher mean active shortening velocity (V_mean_), faster shortening velocity at peak force (V_pkF_), shorter length at the time of 50% force development (L_T50_), and higher mean frequency of EMG activity (E_freq_) (Fig. 5, Fig. S1). These differences are quantified by the ‘treatment’ effect and coefficients in the ANOVA (Tables 1 & 2).

**Figure 5.**
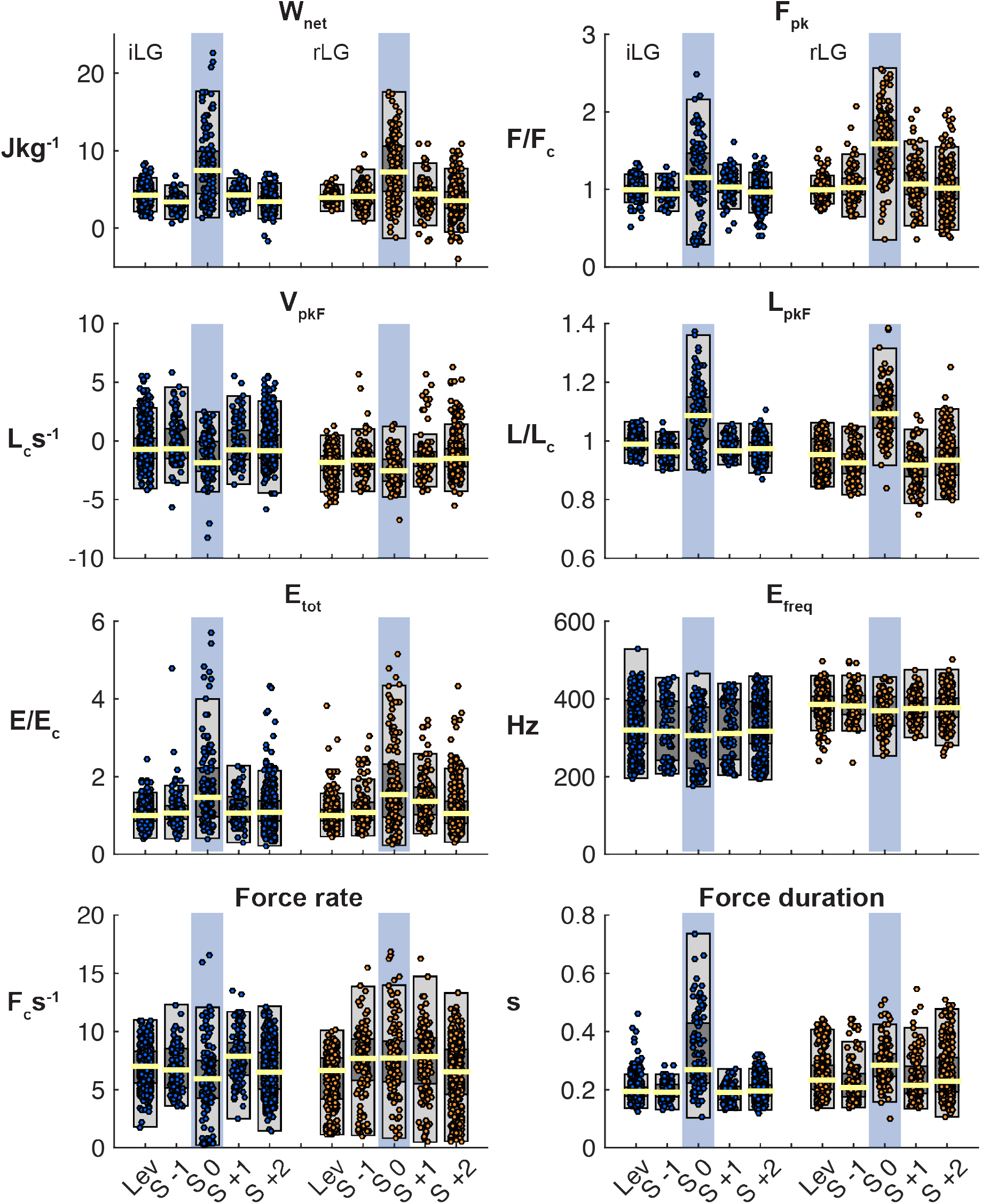
Box plots of median, interquartile range and 1.5 × IQR (yellow line, dark grey box, and light grey box, respectively), overlaid with data points for all measured strides of iLG (blue dots) and rLG (orange dots). Data are separated into stride categories for level and obstacle terrain (Lev, S −1, S 0, S +1, S +2), with the shaded box indicating obstacle contact. Data are shown for total work output (W_net_), peak force (F_pk_), velocity and length at peak force (V_pkF_, L_pkF_), total EMG activity (E_tot_) mean frequency of EMG activity (E_freq_), rate of force development and total force duration. See Tables 1-2 for additional measured variables and statistical results.

Additionally, the steady-state timing of rLG activation (EMG) is phase-shifted to 6% (23ms) earlier in the stride cycle relative to the length trajectory (Fig. 6), quantified by the variable ‘E_phase_’ in the ANOVA (Table 1, Table 2). Earlier onset of activation may help explain the higher rate of shortening throughout stance, with fewer strides exhibiting the stretch or isometric phase that often occurs in early stance in iLG (Fig. 4). Although the total muscle work produced (W_net_) is similar in magnitude between reinnervated and intact conditions, the mechanisms of work production differ (Fig. 5). In iLG increased work upon obstacle contact occurs through increased peak force (F_pk_) and force duration (Fig. 5, S 0, blue). In rLG, increased work occurs through increased F_pk_ and faster shortening velocity (V_pkF_) (Fig. 5, S 0, orange). Force duration does not consistently increase in obstacle strides for rLG but does increase for iLG (Fig. 5).

**Figure. 6.**
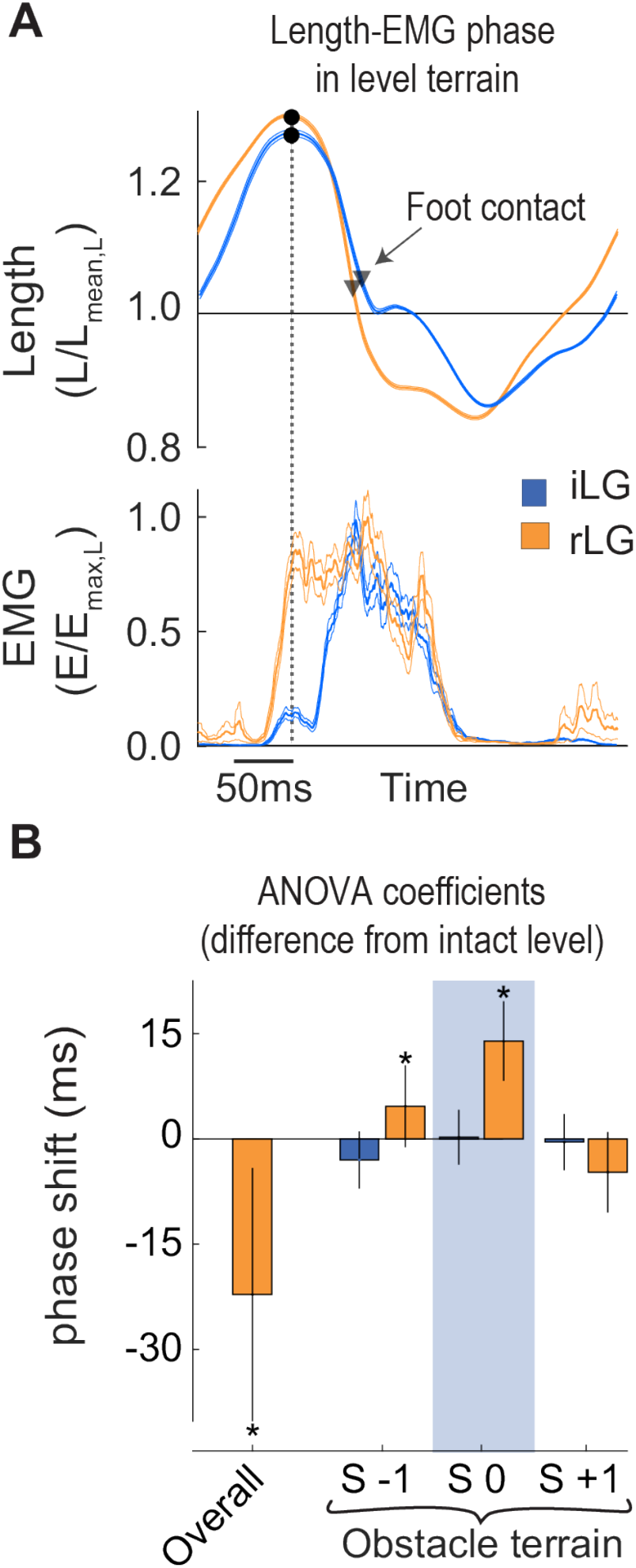
Phase relationship between length and EMG activation (E_phase_) **A)** Average steady-state length and activation trajectories for iLG and rLG in level terrain, aligned in time based on peak length during the swing phase, before foot- substrate contact. Black dot and vertical dashed line indicate the time of peak fascicle length. Timing of foot contact is indicated by downward triangles. **B)** Linear mixed effect model coefficients and 95% confidence intervals for E_phase._ The ‘Overall’ term corresponds to the steady-state difference between iLG and rLG across both level and obstacle terrain. Within obstacle terrain, coefficients are shown for three stride categories: preceding, on and following obstacle contact (S −1, S 0, S +1), with the shaded box indicating obstacle contact. E_phase_ is reported in the ANOVA tables as a fraction of the stride cycle but has been converted to milliseconds here.

### Shifts in timing of obstacle-induced increases in EMG activity between iLG and rLG

To further explore the timing of obstacle-induced changes in LG force, length and activation, we calculated the difference between the average trajectories for steady-state level strides (Lev) and obstacle perturbed strides (S 0) for each individual, and then calculated the mean and 95% confidence interval across individuals for the deviation trajectory (Fig. 7).

**Figure. 7.**
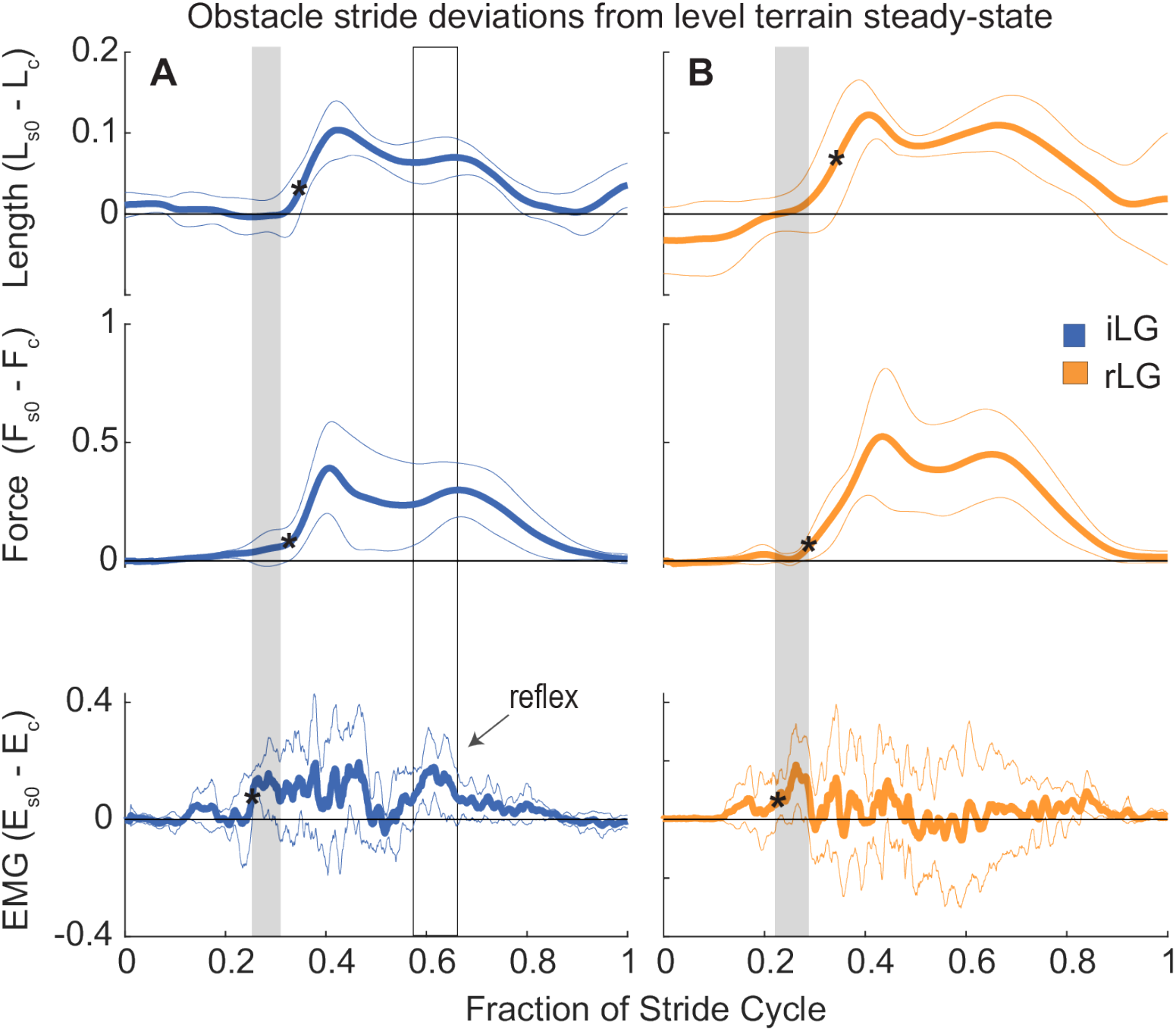
Deviations in the trajectories of LG muscle length, force and activation in obstacle perturbed strides (S 0) compared to steady level terrain (grand mean and 95% CI across individuals). The horizontal zero line indicates no difference from level terrain. Black asterisks (*) indicate the first timepoint that is significantly different from level terrain. The shaded box highlights the time period with increased EMG activity before significant deviations in length and force. This suggests an anticipatory increase in neural drive to the LG. The outlined black box in **A** indicates a 2^nd^ period of increased EMG in late stance in the iLG, suggesting a reflex-mediated response. In **B**, the wide 95% confidence intervals for rLG EMG in late stance indicate inconsistent patterns of activity across individuals, despite consistent deviations in force and length. This suggests disrupted autogenic proprioceptive reflexes and idiosyncratic heterogenic feedback patterns across individuals.

Although the average total increase in E_tot_ is similar (Table 2), the timing of increased activation in obstacle encounters differs between iLG and rLG. Increased neural activation begins 30-40 ms *before* changes in length and force in obstacle strides, for both iLG and rLG, (Fig. 7), suggesting an anticipatory (feedforward) contribution. In iLG, the anticipatory increase in EMG starts ~25% of stride period, which is followed by a later burst of increased EMG activity starting ~58% of stride period, suggesting both feedforward and reflex-mediated feedback contributions to increased EMG activity in obstacle contact strides (Fig. 7). In rLG, the anticipatory increase in EMG starts around 21% of stride period, however activity in the latter half of stance is highly variable and idiosyncratic among individuals, as indicated by the wide 95% confidence intervals spanning −0.2 to +0.2 of level EMG intensity for the period between 40-75% of the stride (Fig. 7, Fig. S2). In contrast, the two distinct bursts of increased EMG in iLG shows greater consistency in the feedforward and feedback contributions across individuals (Fig. 7). Cross-correlation between the obstacle perturbation trajectories for iLG reveals a correlation of 0.82 between length and EMG deviations, and a correlation of 0.85 between force and EMG deviations. For rLG, the cross-correlations are reduced to 0.53 between length and EMG deviations, and 0.57 between force and EMG deviations, respectively. The reduced correlations suggest a disrupted reflexmediated response to muscle load and strain in the latter half of stance in rLG, but which is present in iLG (Fig. 7). Considering that the changes in muscle length and force are strongly correlated with each other during obstacle encounters in both intact and reinnervated conditions (Fig. 7), it is difficult to distinguish the specific sensory signal eliciting reflex responses.

### Stability and kinematic changes during obstacle negotiation in intact vs reinnervated birds

Several differences between iLG and rLG suggest that birds with rLG have reduced stability in obstacle terrain. Differences in the obstacle response and recovery are revealed by the interaction term (treatment × stride category) in the linear mixed effect ANOVA (Tables 1-2). Obstacle-induced (S 0) increases in peak force (F_pk_), force impulse (J_tot_) and work output (W_net_) are higher in magnitude and variance for rLG compared to iLG (Fig. 5, Table 2). In rLG, a larger number of significant differences from level terrain persist in the stride following obstacle contact (S +1), as indicated by significant increases in F_pk,_ J_tot,_ V_pk_ and E_tot_ in rLG but not iLG (see ‘Tr × (S +1)’ coefficients in Table 2). Additionally, rLG shows increases in peak force (F_pk_) and force impulse (J_tot_) in strides preceding obstacle contact (S −1) and in non-obstacle strides within obstacle terrain (S +2), but with higher variance compared to iLG (Fig. 5, Table 2). In contrast, iLG shows lower force and work output in non-obstacle strides within the obstacle terrain, and similar variance as in level terrain (Fig. 5, Table 2). Muscle activation (E_tot_) increases by 11-12% on average for both rLG and iLG in non-obstacle strides within obstacle terrain, compared to level terrain (Table. 2). In rLG there is also an additional 27% increase in E_tot_ increase in S +1, immediately following obstacle contact. Taken together, these results suggest that both iLG and rLG face increased activation costs to maintain stability in obstacle terrain, but the rLG exhibits higher variance and slower recovery to steady state patterns of activation, force and length dynamics, indicating reduced stability.

Reinnervated birds also exhibited differences in running kinematics in obstacle terrain, undergoing a more pronounced ankle flexion in obstacle encounters (Fig. 8), and showing higher variation in stride duration, with a shorter stride in preceding the obstacle encounter (S −1), which is not observed in intact birds (Fig. 8). These kinematic changes are consistent with greater feedforward preparation for the obstacle encounter in reinnervated birds.

**Figure. 8.**
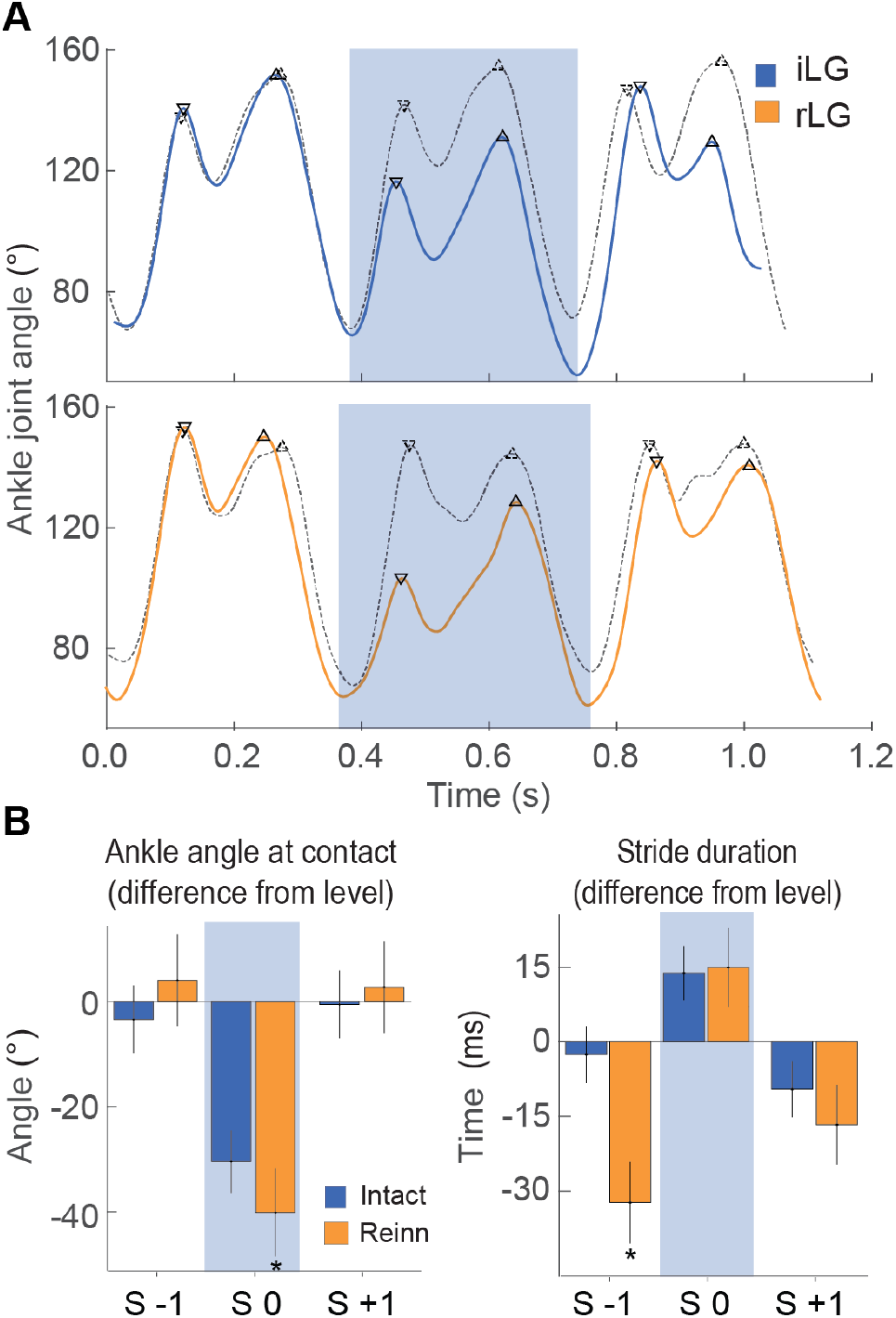
Ankle kinematics in guinea fowl with intact and reinnervated lateral gastrocnemius (LG). **A)** Example ankle joint angle trajectories for a bird with intact LG (blue, top) and a bird with reinnervated LG (orange, below), running over level and obstacle terrain (dashed and solid lines, respectively). **B)** Linear mixed effect ANOVA coefficients and 95% confidence intervals for three obstacle terrain stride categories (S −1, S 0, S +1) compared to level terrain. In obstacle strides (S 0, shaded box), the ankle is more flexed at foot contact in reinnervated birds compared to intact birds. Reinnervated birds show a shorter stride period in S −1, preceding the obstacle encounter, suggesting anticipatory preparation.

## Discussion

### What is the role of proprioception in the control of high-speed locomotion?

We investigated the role of proprioceptive feedback in the sensorimotor control of running by examining the effects of chronic, bilateral autogenic proprioception deficit in the lateral gastrocnemius muscle (LG) of guinea fowl. Measurement of *in vivo* muscle dynamics in a bipedal animal provides opportunity to probe the mechanisms of sensorimotor integration and plasticity that enable stable bipedal gait. Long sensorimotor delays relative to limb cycling times necessitate that animals use a combination of feedforward, feedback and intrinsic mechanical control mechanisms to achieve stable locomotion at high speeds (Brown & Loeb, 2000; Daley & Biewener, 2011; Daley et al., 2009; Frigon & Rossignol, 2006; Grillner, 2011; Lam & Pearson, 2002; More & Donelan, 2018; Pearson & Gramlich, 2010; Prochazka & Ellaway, 2012). We hypothesized that an autogenic proprioceptive deficit will lead to increased reliance on feedforward tuning of muscle activity to achieve stable muscle dynamics in obstacle terrain. In birds with intact LG proprioception, the timing of muscle activity in obstacle-perturbed strides is consistent with combined feedforward and feedback control (Daley & Biewener 2011, Gordon et al. 2015). Birds with reinnervated LG (rLG) exhibit a consistent phase shift in EMG relative to muscle length, with activation starting 6% earlier (23ms) in the steady state contraction cycle, in both level and obstacle terrain (Fig. 6, Table 2). This is consistent with a feedforward tuning of rLG activation timing to enable rapid force development and higher muscle stiffness at the time of foot contact, in the absence of monosynaptic reflexes. Birds with rLG also show anticipatory changes in stride timing and ankle flexion, consistent with feedforward adjustment to maintain intrinsic stability following loss of proprioception. Regulation of force duration in obstacle strides is absent in rLG (Fig. 5), suggesting that proprioceptive feedback in late stance normally regulates force duration, which is disrupted following reinnervation.

A stable intrinsic mechanical response with neither feedforward- nor feedback-mediated changes in neural drive to the muscle can occur when an unexpected perturbation is encountered at high running speeds (Daley et al. 2009). Rapid changes in muscle length and velocity in response to perturbations can decouple activation and force development (Daley et al. 2009, Daley & Biewener, 2011). In our previous work on intact *in vivo* muscle dynamics, variation in LG muscle strain during the initial foot contact and loading period explained 60% of the variation in total force impulse in obstacle perturbation responses while variation in LG muscle activation explained only 9%. This clearly demonstrates the decoupling between activation and force development that can occur *in vivo* (Daley & Biewener, 2011). These intrinsic mechanical effects minimize the disturbances in body dynamics that arise from terrain height perturbations, enabling recovery to steady gait within 2-3 strides. Similar intrinsically stabilizing responses have been demonstrated in the distal hindlimb joints of hopping and running humans subjected to unexpected changes in terrain height and stiffness (Dick, et al., 2019; Ferris, et al., 1999; Moritz & Farley, 2004). In concert with the stabilizing contributions of the intrinsic muscle-tendon dynamics, guinea fowl with intact proprioception also use both feedforward and feedback regulation of muscle activity to maintain stability in obstacle terrain, with greater feedforward contributions when obstacles are visible and high contrast (Daley & Biewener, 2011, Gordon et al. 2015).

We find that guinea fowl with LG proprioceptive deficit achieve similar total increases in EMG magnitude during obstacle strides, demonstrating that they do not rely on intrinsic mechanics alone following reinnervation. The increases in activity occur early in the stride, before obstacle-induced changes in muscle force and length (Fig. 7); consistent with anticipatory, feedforward increases in neural drive to the muscle, similar to that observed in birds running over high-contrast visible obstacles (Gordon et al. 2015), and humans hopping on randomized but expected increases in surface stiffness (Moritz & Farley, 2004). These findings are consistent with a hybrid feedforward/feedback control model as conceptualized by Kuo (2002) in which feedforward and feedback gains are balanced to enable accurate state estimation and robust cyclical dynamics in the presence of both disturbances and sensory error.

We do observe shifts in muscle activity in late stance in some individuals in response to obstacle encounters, which suggests heterogenic reflex responses in reinnervated LG (Fig. 4). However, these responses are variable and idiosyncratic, with some birds showing increased EMG in late stance in obstacle strides, and others showing a decrease (Fig. S2). The variable and idiosyncratic reflex responses result in wide confidence intervals for the obstacle-induced EMG response in the latter half of stance, despite consistent force-length trajectories over the same time-period (Fig. 7). The idiosyncratic use of heterogenic reflex modulation across individuals following nerve injury recovery is consistent with findings in cats (Lyle & Nichols, 2017). Guinea fowl have several agonist muscles to the LG that could contribute to heterogenic feedback modulation, including the medial gastrocnemius and digital flexors (Daley & Biewener, 2011, Gordon et al. 2015). However, no muscle is an exact synergist of the LG, because each has a unique combination of moment arms, fiber length, pennation angle and connective tissue compliance (Daley & Biewener 2003; Cox et al. 2019). Consequently, it is unlikely that proprioception from agonists can completely restore accurate sensing to regulate LG force and work output. Additionally, in the presence of increased sensory error and noise, birds may learn over time to compensate through sensory integration in higher CNS pathways, leading to updated central coordination and feedforward drive to rhythm generating networks.

Despite variability in rLG EMG activity patterns across individuals, we observe recovery of similar work output and stabilizing responses in obstacle terrain following self-reinnervation. Reinnervated LG achieves qualitatively similar stabilizing response to obstacle encounters, but with multiple differences in the underlying neuromuscular mechanisms (Fig. 9). The shift to earlier rLG activation and more flexed ankle kinematics suggests coordinated plasticity and tuning of feedfoward control and intrinsic mechanics to maintain ankle stiffness and stable locomotion following recovery from nerve injury. Earlier activation enables higher muscle force development to resist the external load applied at foot contact, and therefore likely contributes to a higher rate of shortening throughout stance (Fig. 9), which enables rLG to achieve similar work output as iLG in both level and obstacle terrain, despite the disrupted regulation of force duration. These changes suggest integrated plasticity of neural and musculoskeletal systems during the recovery from nerve injury. The findings also suggest that sensorimotor mechanisms may be optimized over time to maintain higher-level performance objectives such as stable body dynamics. Similarly, cats that have recovered from nerve injury exhibit high variance in inter-joint coordination but preserve whole-limb function (Chang et al. 2009), suggesting that consistent performance of task-level locomotor goals is a target of sensorimotor optimization. Nonetheless, although the rLG achieves an effective stability response, it requires relatively higher activation for a less effective outcome— the reflex response of the intact system enables robust stability with lower energy costs of muscle activation.

**Figure 9.**
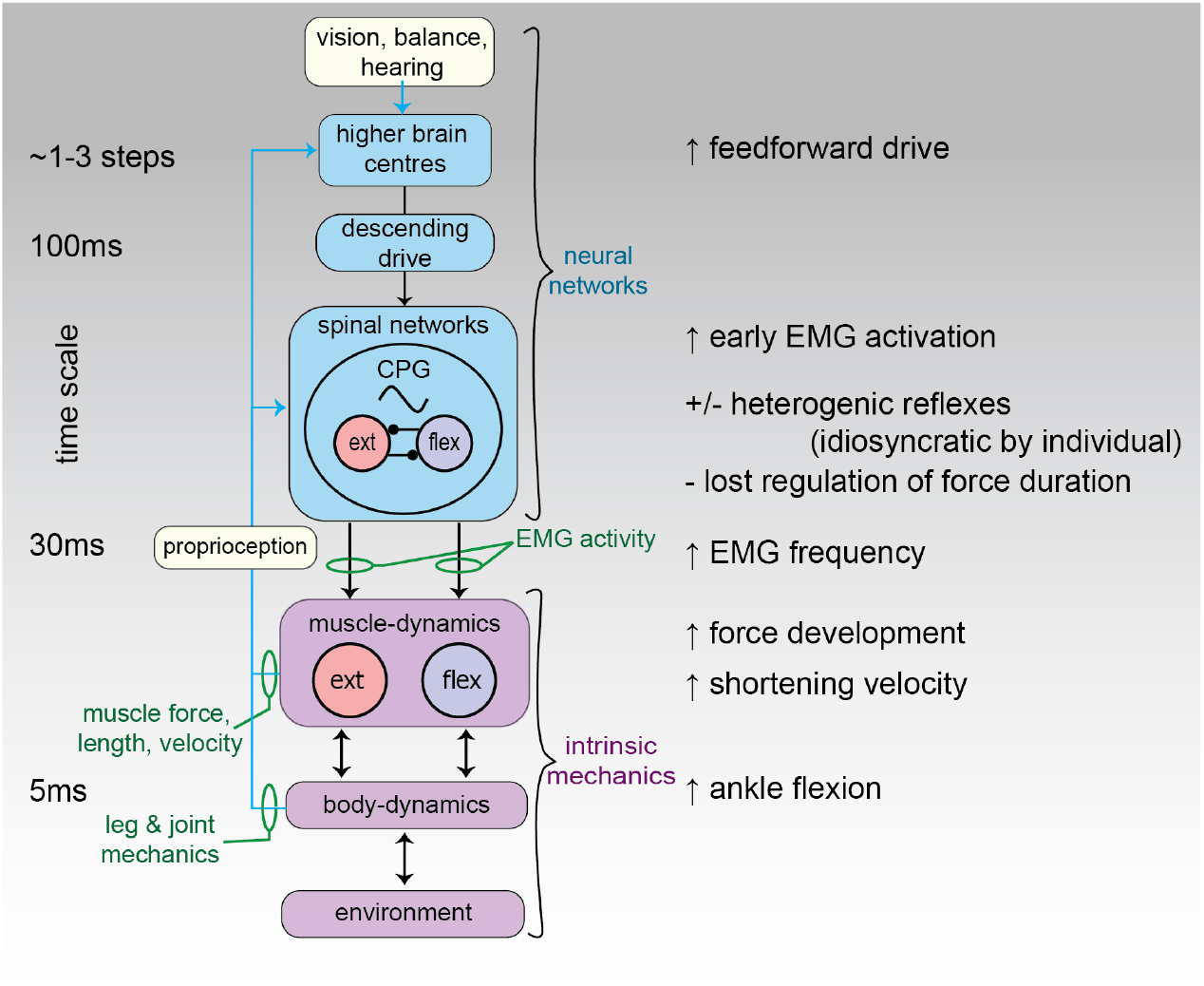
Schematic illustration of the neuromechanical control system for locomotion, indicating the mechanisms observed to change in response to self-reinnervation in the lateral gastrocnemius (LG) of guinea fowl. Green text indicates the *in vivo* experimental measures used in this study to infer sensorimotor control mechanisms. On the right, black text summarizes the changes observed in the control and muscle mechanics of the LG in response to the self-reinnervation recovery and chronic autogenic proprioceptive deficit. Guinea fowl with proprioceptive deficit maintain qualitatively similar force-length muscle dynamics and stabilizing responses to obstacle encounters during running. However, they use a combination of feedforward and intrinsic mechanical control mechanisms to compensate for disrupted proprioceptive reflexes, suggesting interconnected plasticity of neural and musculoskeletal mechanisms in the recovery from nerve injury.

A recent study by Sawicki et al. (2015) found that earlier onset of activation was associated with a shift to energy absorption in cyclical muscle contractions with a sinusoidal MTU length trajectory. We find here that earlier onset is associated with greater shortening and work production. The specific response of a muscle to a shift in activation phase is likely to be highly sensitive to the specific steady-state length trajectory of the muscle. Muscle force capacity and the activation and deactivation kinetics are substantially influenced by velocity and recent strain history, as demonstrated in controlled studies of *in vitro* muscle force-length work loops (Askew & Marsh, 1998; Josephson, 1999). Further work is needed to understand how *in vivo* muscle fascicle length dynamics interact with neural activation patterns and MTU compliance to enable tuning of muscle contraction dynamics to the mechanical demands of cyclical locomotor tasks.

It is important to note that the self-reinnervation procedure requires a long-term recovery period and results in a chronic sensory deficit, which is likely to lead to a complex array of changes in the musculoskeletal tissues and the sensorimotor networks. The recovery process almost certainly involves coupled changes across multiple systems, including connective tissue compliance, muscle activation kinetics, fiber type distribution, motor unit size and distribution, spinal intraneuronal connectivity, and sensory integration in higher CNS centers for state estimation and movement planning. Due to the complex nature of these adaptations, it is challenging to fully tease apart individual contributions and mechanisms from *in vivo* experimental measures alone. Nonetheless, the current results provide empirical evidence for feedforward tuning of muscle activation and ankle kinematics to maintain intrinsic stability following loss of proprioception.

In future studies, these coordinated mechanisms of sensorimotor adaptation and plasticity could be systematically explored through a combination of integrative experimental and computational approaches. These approaches could include 1) closed loop neuromechanical simulations to enable predictive hypothesis testing (Ijspeert, 2014; Roth, et al., 2014), 2) combined use of *in vivo* measures of muscle dynamics with *in vitro* testing of muscle contractile dynamics, to replicate biologically realistic force-length contraction dynamics, 3) histological studies to examine changes in muscle fiber type distribution and connective tissue characteristics following reinnervation, and 4) perturbation approaches that probe both short and long term adaptation processes.

It remains unclear how the specific length trajectory and velocity features of *in vivo* muscle dynamics contribute to the intrinsic stability and control of movement. Dynamic measurement techniques are needed to address this challenge and to develop realistic models for *in vivo* muscle-tendon function. In additional to widely recognized force-length and force-velocity ‘Hill-type’ properties, muscle exhibits well-documented short and long-term history-dependent changes in force capacity in response to stretch and shortening (Edman, 1975; Edman et al. 1978; Edman, 1980; Josephson, 1999; Herzog, 2004; Edman 2012; Herzog 2014; Rode et al. 2009; Nishikawa et al. 2012; Yeo et al. 2013; Nishikawa, et al., 2018). Recent developments in biorobotic platforms that enable controlled muscle experiments with realistic loading and length trajectories are promising tools for advancing our understanding of the role of intrinsic muscle dynamics in the control of movement (Clemente & Richards, 2012; Richards, 2011; Robertson & Sawicki, 2015). Integrative neuromechanical studies using multiple techniques will be essential for unravelling mechanisms of muscle function, sensorimotor integration and plasticity, and findings from these studies have important implications for many human health conditions, including acute nerve injury, diabetic neuropathy, neurodegenerative disorders, cerebral palsy, and muscular dystrophies.

## Acknowledgements

This research was supported by NIH grant NIAMS 5R01AR055648 to AAB, grant BB/H005838/1 to MAD from the Biotechnology and Biological Sciences Research Council (BBSRC), and a doctoral training studentship from the BBSRC to JCG supervised by MAD. Thanks to Jennifer A Carr for assistance in the experiments.

## Author Contributions

AAB and MAD conceived the study, JCG, AAB and MAD designed the experiments, JCG and AAB conducted surgical procedures, JCG, AAB, MAD and NCH collected experimental data, JCG and MAD analyzed the data and wrote the paper. All authors commented on drafts and approved the manuscript.

## Supplemental Figures

**Figure S1.**
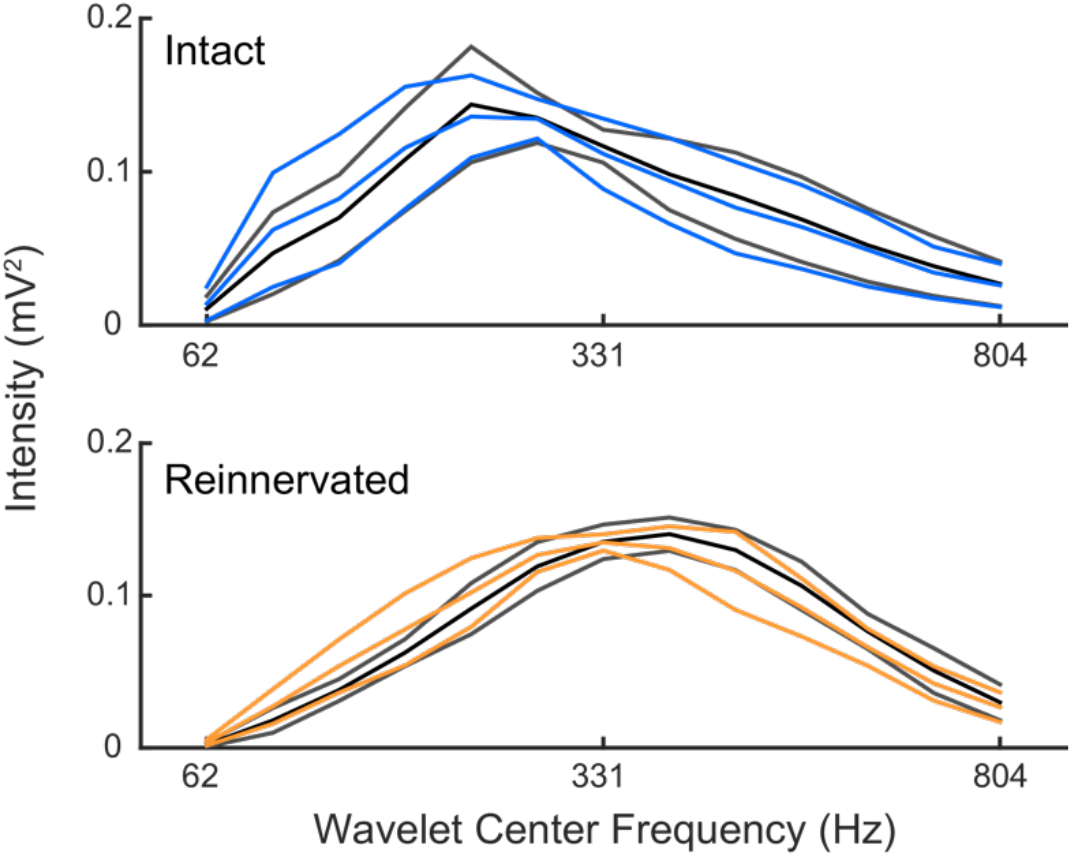
Distribution of EMG activation frequency (mean ± 95% confidence interval) for intact LG (top, blue) and reinnervated LG (bottom, orange), during level running (dark grey lines) and obstacle encounters (S 0, colored lines). Note the shift in peak frequency of EMG activity in the reinnervated LG for both level terrain and obstacle strides, suggesting recruitment of faster motor units.

**Figure S2.**
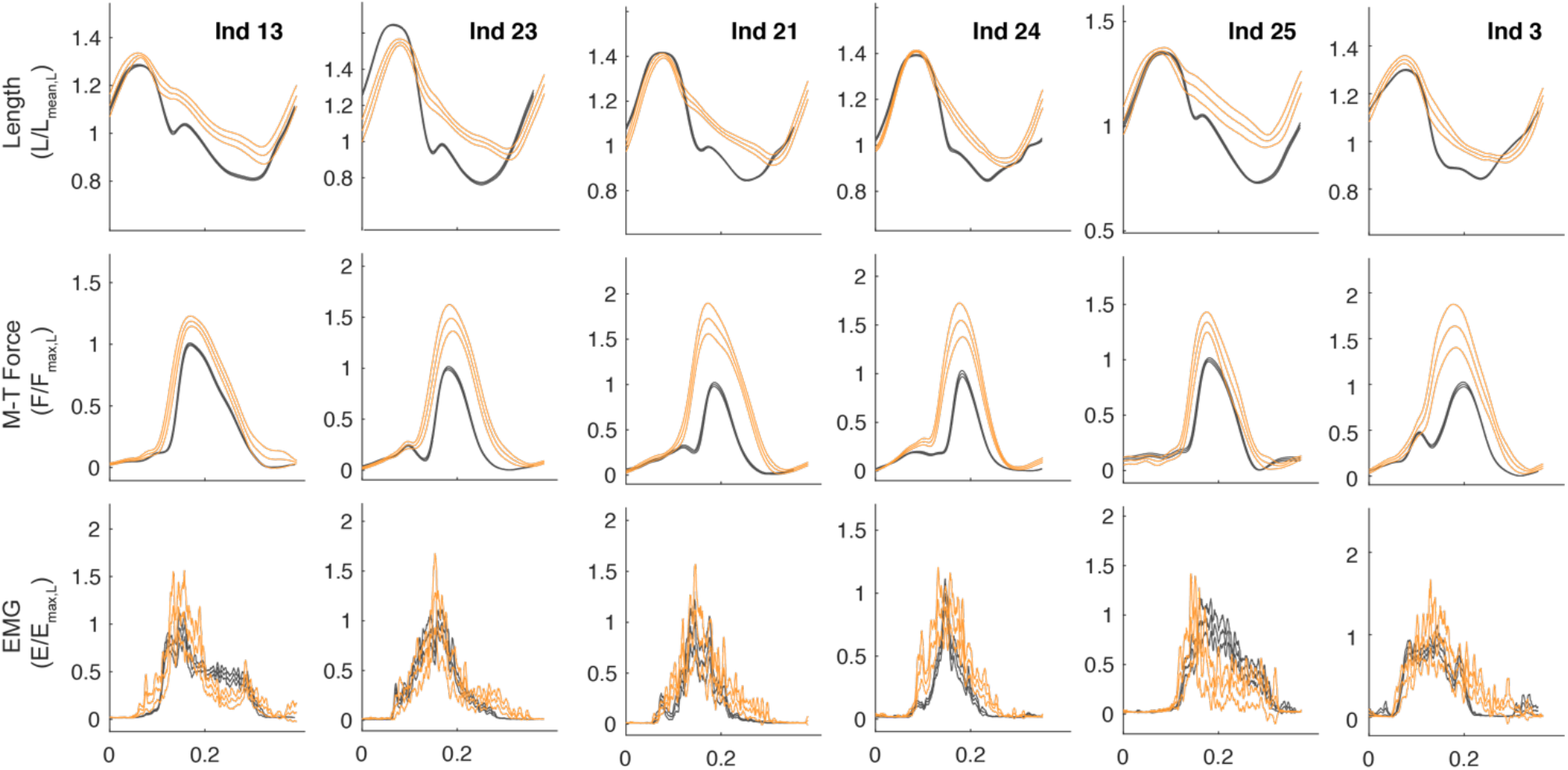
Average stride cycle trajectories (mean± 95% confidence intervals) for fascicle length, muscle-tendon force and myoelectric activity (EMG) during obstacle encounter strides (S 0, orange) compared to the level terrain averaged (grey), for all birds with reinnervated LG. Although the deviations in length and force during obstacle encounters is qualitatively similar across individuals, the shifts in EMG activation in the latter half of stance varies substantially across individuals, with some individuals showing reflex inhibition of activity (Ind 13, Ind 25) and others showing reflex excitation.

